# Matching the forecast horizon with the relevant spatial and temporal processes and data sources

**DOI:** 10.1101/807057

**Authors:** Peter B. Adler, Ethan P. White, Michael H. Cortez

**Affiliations:** Department of Wildland Resources and the Ecology Center, Utah State University, Logan, Utah; Department of Wildlife Ecology and Conservation, University of Florida, Gainesville, Florida; Informatics Institute, University of Florida, Gainesville, Florida; Biodiversity Institute, University of Florida, Gainesville, Florida; Department of Biological Science, Florida State University, Tallahasee, Florida

**Keywords:** dispersal, ecological forecasting, eco-evolutionary dynamics, global change, selection

## Abstract

Most phenomenological, statistical models used to generate ecological forecasts take either a time-series approach, based on long-term data from one location, or a space-for-time approach, based on data describing spatial patterns across environmental gradients. Here we consider how the forecast horizon determines whether more accurate predictions come from the time-series approach, the space-for-time approach, or a combination of the two. We use two simulated case studies to show that forecasts for short and long forecast horizons need to focus on different ecological processes, which are reflected in different kinds of data. In the short-term, dynamics reflect initial conditions and fast processes such as birth and death, and the time-series approach makes the best predictions. In the long-term, dynamics reflect the additional influence of slower processes such as evolutionary and ecological selection, colonization and extinction, which the space-for-time approach can effectively capture. At intermediate time-scales, a weighted average of the two approaches shows promise. However, making this weighted model operational will require new research to predict the rate at which slow processes begin to influence dynamics.

## Introduction

Forecasting is increasingly recognized as important to the application and advancement of ecological research. Forecasts are necessary to guide environmental policy and management decisions about mitigation and adaption to global change (Clark et al., 2001; Mouquet et al., 2015; Dietze et al., 2018). But forecasts can also advance understanding of the processes governing ecological systems by providing rigorous tests of model predictions (Houlahan et al., 2017; Dietze, 2017; Dietze et al., 2018). The dual benefits of informing management and advancing basic knowledge makes forecasting an important priority for ecological research.

Statistical models used for ecological forecasting generally rely on either time-series approaches or space-for-time substitutions. The time-series approach involves fitting models to long-term datasets to describe the temporal dynamics of a system. We then use those dynamic models to make predictions about what will happen in the future. This approach is often used to study population or vital rate fluctuations as a function of weather (Dalgleish et al., 2011), or primary production as a function of annual precipitation (Lauenroth and Sala, 1992). When time-series models are fit with typically short ecological data sets, they capture “fast processes” operating on interannual time-scales, such as birth, death, individual growth, small-scale dispersal events, and short-term responses to environmental conditions (Fig. 1). Statistical models built using this approach normally cover a limited spatial extent (but see Hefley et al. 2017; Kleinhesselink and Adler 2018; Chevalier and Knape 2020), and ignore slower processes, such as evolutionary adaptation or turnover in community composition, that could influence dynamics at longer time scales (Clark et al., 2001).

**Figure 1:**
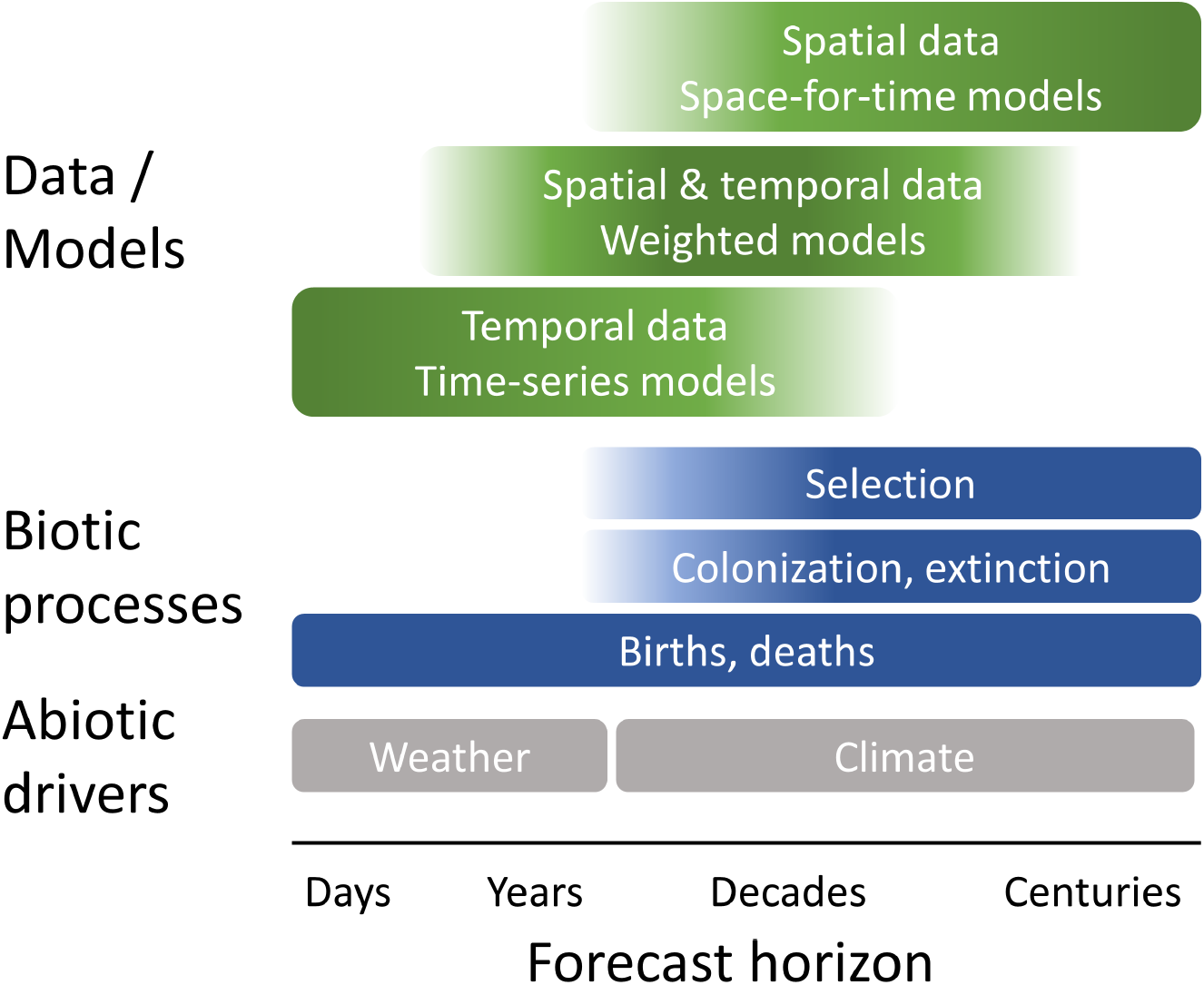
Fast and slow processes operate at different time scales, and are reflected in different kinds of datasets. Fast processes, such as births, deaths, and individual growth, operate at all time scales, but are the exclusive drivers of the short-term dynamics captured in most time series datasets. Slower processes, such as evolutionary selection on genotype frequencies, ecological selection on species abundances, and colonization and extinction, interact with fast processes to drive dynamics over the long-term. The influence of these slow processes is seen in very long time series, or in spatial gradients. Understanding dynamics at intermediate time scales requires integrating information from spatial and temporal data sources. We propose a model weighting approach; mechanistic spatiotemporal modeling is another alternative. The time scales shown here were chosen with vascular plants in mind, but the same concepts would apply for much shorter-lived organisms but at shorter time scales.

Space-for-time substitution approaches begin by describing how an ecological variable of interest, such as occupancy or productivity, varies across sites experiencing different environmental conditions. These spatial relationships between environment and ecological response are assumed to also hold for changes at a site through time. To make a forecast, we first predict the future environmental conditions and then determine the associated ecological response, based on the observed spatial relationship. This is the approach commonly used to predict population distribution or abundance as a function of climate (Elith and Leathwick, 2009) or mean primary production as a function of mean precipitation (Sala et al., 1988). Space-for-time models capture the outcome of interactions between fast processes and slower processes operating over long time periods, such as immigration, extinction, and responses to large or prolonged environmental changes (Fig. 1). However, space-for-time models provide no information about how quickly the system will move from the current state to the predicted, future state. In fact, transient dynamics could prevent the system from ever reaching the predicted steady state (Urban et al., 2012). Although both time-series and space-for-time approaches are widely used, there has been little discussion of their advantages and disadvantages for guiding policy decisions or advancing our understanding of ecological dynamics (Harris et al., 2018; Renwick et al., 2018).

Whether historical dynamics, contemporary spatial patterns, or some combination of the two will serve as the best source of information for forecasting may depend on how far into the future we are attempting to forecast (Harris et al., 2018). This potential dependency on the “forecast horizon” (*sensu* Hyndman and Athanasopoulos 2018) reflects lags in the response of ecological conditions to environmental change, shifts in the importance of ecological processes with time scale (Levin, 1992; Rosenzweig et al., 1995), and differences between time-series and spatial gradients in the range of environmental conditions represented in observed data (Fig. 1). At short forecast horizons (days to years), dynamics will reflect physiological and demographic responses and interactions among the organisms present at a site more than temporal turnover of genotypes or species; environmental conditions are likely to stay within the range of historical variation; and the current state of the system is likely to capture the influence of unmeasured processes. As a result, for near-term forecasts time-series approaches may capture the key dynamics and provide accurate predictions.

In contrast, at long forecast horizons (decades to centuries), environmental conditions that have not been historically observed are likely to not only occur but to persist long enough to drive significant turnover of genotypes and species through colonization and extinction as well as changes in the flux of energy and nutrients. At these long forecast horizons, the state of the system at the time the forecast is issued may be little help in predicting the future state. For the century-scale forecasts often featured in biodiversity and species-distribution modeling, space-for-time approaches may effectively capture the response of ecosystems to major shifts in climate over long periods, producing better long-term forecasts than time-series approaches. Using different modeling approaches for different forecast horizons is common in other disciplines. For example, meteorological models for short-term weather forecasts differ substantially in spatial and temporal resolution and extent from the global circulation models used to predict long-term changes in climate.

Why not simply use process-based models to avoid the difficulties posed by phenomenological time-series and space-for-time modeling approaches? If we could accurately characterize all of the processes governing a system, then a model based on that understanding should make accurate predictions at all forecast horizons. Process-based models should also be more robust for making predictions outside of historically observed conditions and even beyond the conditions observed across spatial gradients, which will be especially important in a future with increasingly novel combinations of environment and species interactions (Williams and Jackson, 2007). Unfortunately, in most cases this approach is not currently feasible because we lack a detailed knowledge of all the complex and interacting processes influencing the dynamics of real ecological systems. Even if the general form of the models were known, estimating the high number of parameters and quantifying how they vary across ecosystems typically requires more data than is currently available even for well-studied systems. Furthermore, the high complexity and corresponding parameter uncertainty of such models can increase predictive errors; simpler time-series models may actually perform better (Ward et al., 2014), though spatial replication can reduce the cost of complexity (Chevalier and Knape, 2020). As a result, models used for ecological forecasting will include at least some phenomenological components. But that does not mean that phenomenological forecast models cannot benefit from process-based understanding. Even if process-level understanding does not enable a fully mechanistic model, it can improve the specification of phenomenological models. Our hypothesis is that different processes may be relevant for different forecast horizons, and that we can act on this knowledge by fitting models to different kinds of datasets.

Here we use two simulated case studies to 1) demonstrate that the best model-building approaches for ecological forecasting depend on the time horizon of the forecast, and 2) explore how time-series and space-for-time approaches might be combined via weighted averaging to make better forecasts at intermediate time scales. The first case study focuses on how interspecific interactions affect the population dynamics of a focal species, and the second focuses on an eco-evolutionary scenario. Our analyses show that:

1. For short-term forecasts, phenomenological time-series approaches are hard to beat, whereas longer-term forecasts require accounting for the influence of slow processes such as evolutionary and ecological selection as well as dispersal.
2. Different kinds of data reflect the operation of different processes: longitudinal data capture autocorrelation and fast responses of current assemblages to interannual environmental variation, while data spanning spatial gradients capture the long-term outcome of interactions between fast and slow processes. Whether predictive models should be trained using longitudinal or spatial data sets, or both, depends on the time-scale of the desired forecast.
3. A key challenge for future research is determining the rate at which slow processes begin to influence dynamics.

## Modeling approach

In each case study, we simulated the effects of an increase in temperature on simple systems with known dynamics. The truth was represented by a simulation model that was mechanistic for at least one important process, but we treated the data-generating model as unknown when analyzing the data and we assumed that perfectly recovering the mechanisms it contains would not be possible in practice. We began each simulation under a stationary distribution of annual temperatures, allowing the system to equilibrate; we call this the baseline phase. We then increased temperature progressively over a period of time, followed by a second period of stationary, now elevated, temperature. The objective was to forecast the response of the system to the temperature increase based on spatial and/or temporal data “sampled” from the simulation during the baseline period.

We made forecasts based on two phenomenological statistical models, each representing processes operating at different time scales. One statistical model represents the time-series or “temporal approach.” We correlated interannual variation in an ecological response with interannual variation in temperature at just one site. The other statistical model relies on a space-for-time substitution, which we call the “spatial approach” for brevity. We correlated the mean temperature with the mean of an ecological state or rate across many sites. We compared forecasts from both statistical models to the simulated dynamics to determine how well the two approaches performed at different forecast horizons. We also assessed the potential for combining the information available in temporal and spatial patterns by using a weighted average of the forecasts from the temporal and spatial approaches optimized to best match the (simulated) observations. We then studied how the optimal model weights changed over time. We expected the temporal approach to best predict short-term dynamics, the spatial approach to best predict long-term dynamics, while the weighted model would show potential to provide the best forecasts at transitional, intermediate time scales. The three statistical models are described in Supporting Information (Appendix A). Computer code for both case studies is available for reviewers online (https://drive.google.com/open?id=1aju04qtQvJmZG1mcpN-6hYBZILBhDfl2) and will be archived at Zenodo upon acceptance.

## Community turnover example

Conservation biologists and natural resource managers often need to anticipate the impact of environmental change on the abundance of endangered species, biological invaders, and harvested species. Although the managers may be primarily interested in just one focal species, skillful prediction might require considering interactions with many other species, greatly complicating the problem. But at what forecast horizon do altered species interactions become impossible to ignore? We explored this question using a metacommunity simulation model developed by Alexander et al. (2018) to study how community responses to increasing temperature depend on the interplay between within-site demography and competitive interactions and the movement of species across sites. The model features Lotka-Volterra competitive interactions among plants within sites that are arrayed along an elevation and temperature gradient. Composition varies along the gradient because of a trade-off between growth rate and cold tolerance: cold sites are dominated by slow-growing species that can tolerate low temperatures, while warm sites are dominated by fast-growing species that are cold intolerant. Multiple species can coexist within sites because all species experience stronger competition from conspecifics than from heterospecifics. Sites are linked by dispersal: a specified fraction of each species’ offspring leaves the site where they were produced and reaches all other sites with equal probability. We provide a more detailed description of the simulation model in SI Appendix B.

We first simulated a baseline period with variable but stationary temperature, followed by a period of rapid temperature increase, and then a final period of stationary temperature. Interannual variation in temperature is the same at all sites, but mean temperature varies among sites. All sites experienced the same absolute increase in mean temperature. We focused on the biomass dynamics of one focal species that dominated the central site during the baseline period.

During the baseline period there were strong spatial patterns across the mean temperature gradient. Individual species, including our focal species, showed classic, unimodal “Whittaker” patterns of abundances across the gradient (Fig. 2A). These spatial patterns are the basis for our spatial statistical model of the temperature-biomass relationship for our focal species (Fig. 2A). In contrast to the strong spatial patterns, population and community responses to interannual variation in temperature within sites were weak. At our focal site in the center of the gradient, the biomass of the focal species was quite insensitive to interannnual variation in temperature, but showed strong temporal autocorrelation (Fig. 2B). Our temporal statistical model estimates this weak, linear temperature effect, along with the strong lag effect of biomass in the previous year.

**Figure 2:**
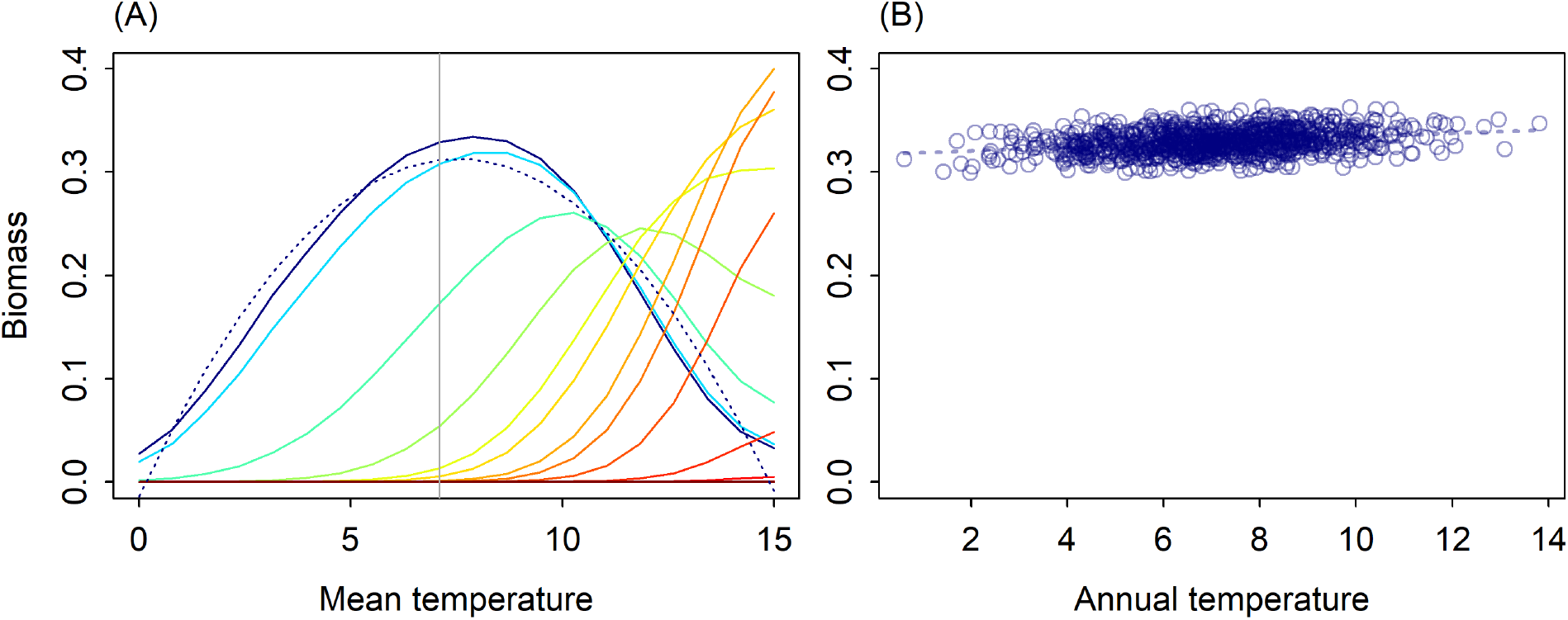
(A) Mean biomass by species (colors) across the temperature gradient during the baseline period. The focal species, dominant at the site in the center of the gradient (vertical gray line), is shown in dark blue. The dashed blue line shows predictions from the spatial statistical model. (B) Annual biomass of the focal species at the central site during the baseline period. The dashed line shows predictions from the temporal statistical model.

We used both the temporal and spatial statistical models to forecast the effect of a temperature increase (Fig. 3A) on the focal species’ biomass at one location in the center of the temperature gradient. The predictions from these two models contrasted markedly, with the temporal statistical model predicting a large increase in biomass and the spatial statistical model predicting a decrease. Initially, the simulated abundances followed the increase predicted by the temporal model, but as faster-growing species colonized and increased in abundance at the focal site, the biomass of the focal species decreased, eventually falling below its baseline level (Fig. 3B).

**Figure 3:**
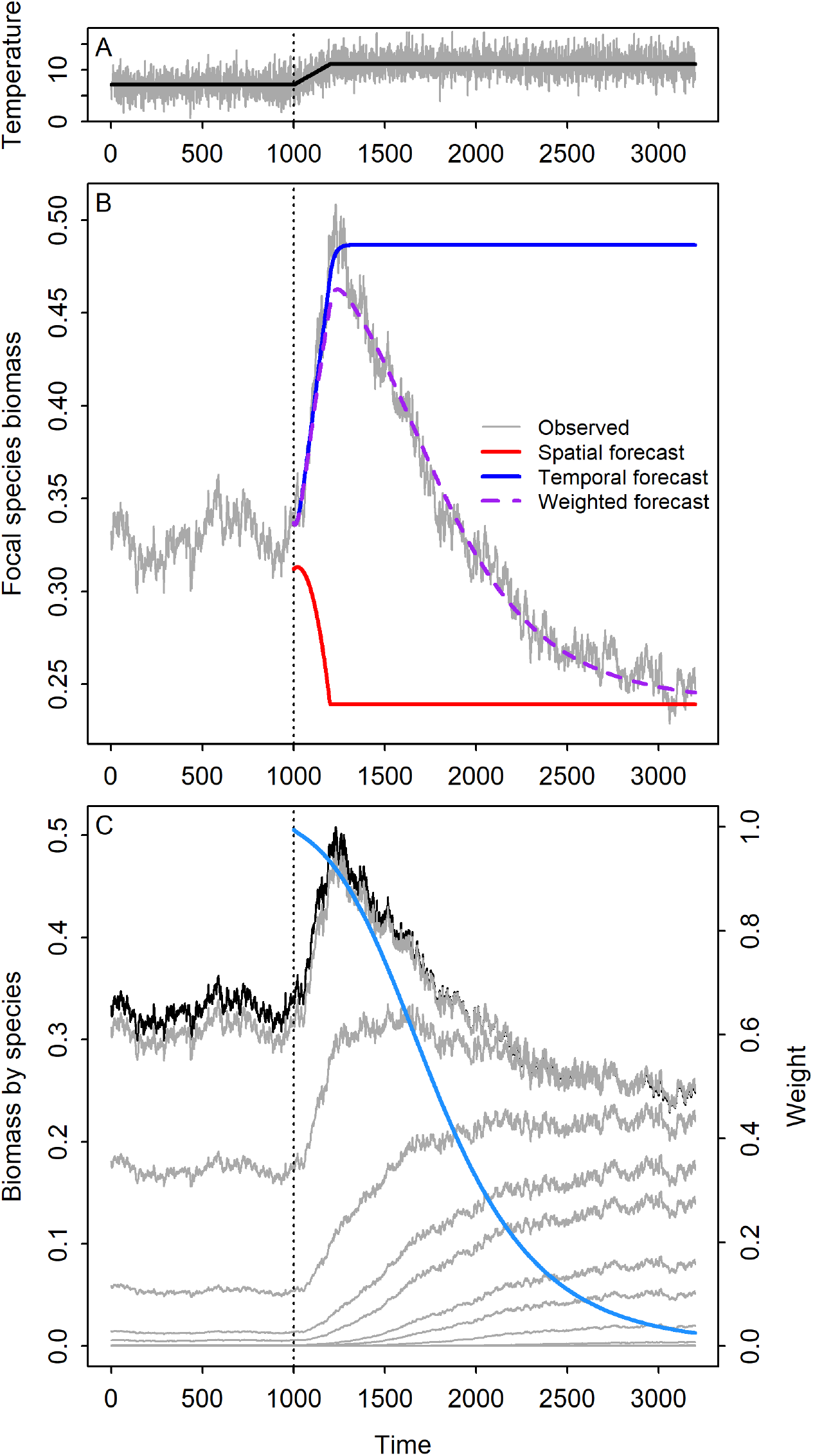
(A) Simulated annual temperatures (grey) and expected temperature (black), which was used to make forecasts, at the focal site. (B) Simulated focal species biomass and forecasts from the spatial, temporal and weighted statistical models at the focal site in the metacommunity model. (C) Simulated changes in biomass of the focal species (black) and all other species (grey), and the weight given to the temporal statistical model for focal species biomass (blue). Year 1000 in each panel corresponds to the start of the temperature increase.

To combine information from the temporal and spatial statistical models into a single prediction, we fit a weighting parameter, *ω*, which varies over time and is bounded between 0 and 1. At any time point, *t*, this weighted forecast is *ω* · *T*(*N*_*t*–1_, *K_t_*) + (1 – *ω*) · *S*(*K_t_*) where *T* is the temporal statistical model, which depends on population size, *N*, and expected temperature, *K*, and *S* is the spatial statistical model, which depends only on *K* (see SI Appendix A for a full description of the approach). The weighted model accurately predicts the simulated dynamics across the full forecast horizon (Fig. 3B). It also shows that the most rapid shifts in the model weights occurred during the period when warm-adapted, faster growing species were increasing most rapidly in abundance (Fig. 3C). However, the reason the weighted models works so well is that the weights were determined by fitting directly to the data. Unlike the forecasts from the spatial and temporal statistical models, we did not generate out-of-sample predictions from the weighted model; it merely provides a convenient way to quantify how rapidly dynamics shift from being dominated by interannual variation captured in the temporal model (time *t* = 0 to *t* ≈ 1250 in Fig. 3B) to being dominated by the steady-state equilibrium captured by the spatial model (time *t* ≥ 2500). A true forecast from the weighted model would require a method to determine the model weights *a priori*.

The compositional turnover affecting our focal species also influences total biomass, linking community and ecosystem dynamics. We repeated our focal species analysis for total community biomass, and the results were similar: the temporal statistical model initially made the best forecasts immediately following the onset of the temperature increase, but as the identity and abundances of species at the study site changed, the model weights rapidly shifted to the spatial statistical model (SI Figs. S-1 and S-2).

## Eco-evolutionary example

Evolutionary adaptation is a key uncertainty in predicting how environmental change will impact a focal population at a given location (Hoffmann and Sgro, 2011). Like the shifts in species composition illustrated in the previous example, shifts in genotype frequencies can also influence dynamics and forecasts at different time scales. Although shifts in genotype frequencies at the population level are analogous to changes in species composition at the community level, the mechanisms are distinct: heterozygosity and genetic recombination have no analogue at the community level. We demonstrate how these processes influence short and long-term forecasts with a standard eco-evolutionary simulation model for a hypothetical annual plant population in which fecundity is temperature dependent, and different genotypes have different temperature optima (Fig. 4A).

**Figure 4:**
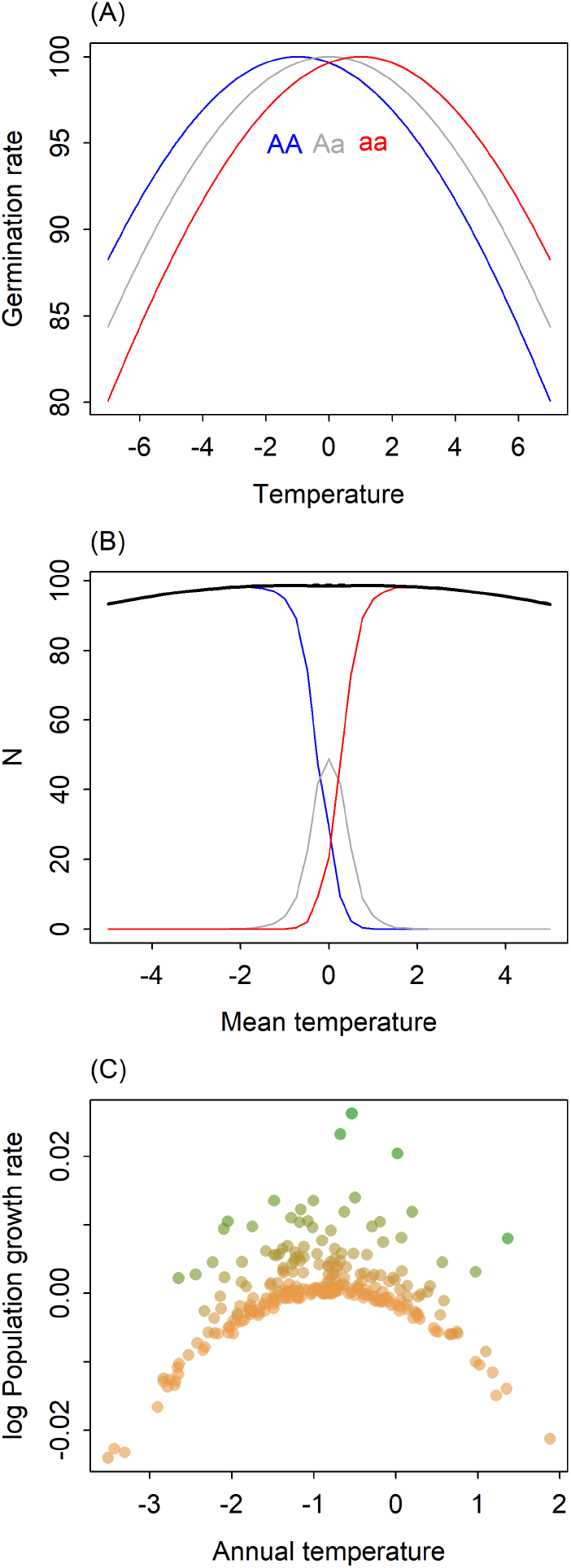
(A) Reaction norms of the three genotypes. (B) The spatial pattern of individual genotypes (colors) and total population abundance (black) at sites arrayed across a gradient of mean annual temperature. The dashed black line (almost entirely hidden by the slid black line) shows predictions from an empirical, spatial statistical model, a linear regression that describes mean population size as a function of mean temperature. (C) The relationship between annual temperature and per capita growth rate at a location with a mean temperature that favors the cold-adapted genotype. Colors show population size (the green to brown gradient depicting low to high population density), which influences the population growth rate through density dependence.

Our model describes how the local density of each genotype changes between years, which depends on temperature and genotype densities in the previous year. Transient temporal dynamics are computed directly from the model; these dynamics are the basis for the temporal statistical model. To create a spatial gradient, we simulated the equilibrium density of each genotype in a series of local populations experiencing different mean temperatures. The pattern of equilibrium densities across the mean annual temperature gradient is the basis for our spatial statistical model: cold sites will be dominated by the cold-adapted homozygous genotype, warm sites will be dominated by the heat-adapted homozygous genotype, and intermediate sites will be dominated by the heterozygous genotype (Fig. 4B). The full description of the eco-evolutionary simulation model is provided in SI Appendix C.

The spatial pattern shown in Fig. 4B is the outcome of steady-state conditions. But at any one site, the population’s short-term response to temperature will be determined by the dominant genotype’s reaction norm (Fig. 4A). For example, at a cold site dominated by the cold-adapted homozygous genotype, a warmer than average year would cause a decrease in population size due to decreases in fecundity (blue line in Fig. 4A), even though the heat-adapted homozygote might perform optimally at that temperature. However, if warmer than normal conditions persist for many years, then genotype frequencies should shift, and the heat-adapted homozygote will compensate for the decreases of the cold-adapted genotype.

To demonstrate these dynamics, we simulated a diploid annual plant population at a colder than average site. During the baseline period, the population is dominated by the cold-adapted genotype. We used the simulated data from this baseline period to fit a temporal statistical model (Appendix A) that predicts population growth rate as a function of annual temperature and population size (Fig. 4C), assuming no knowledge of the underlying eco-evolutionary dynamics. We then imposed a period of warming, followed by a final period of higher stationary temperature (Fig. 5 top).

**Figure 5:**
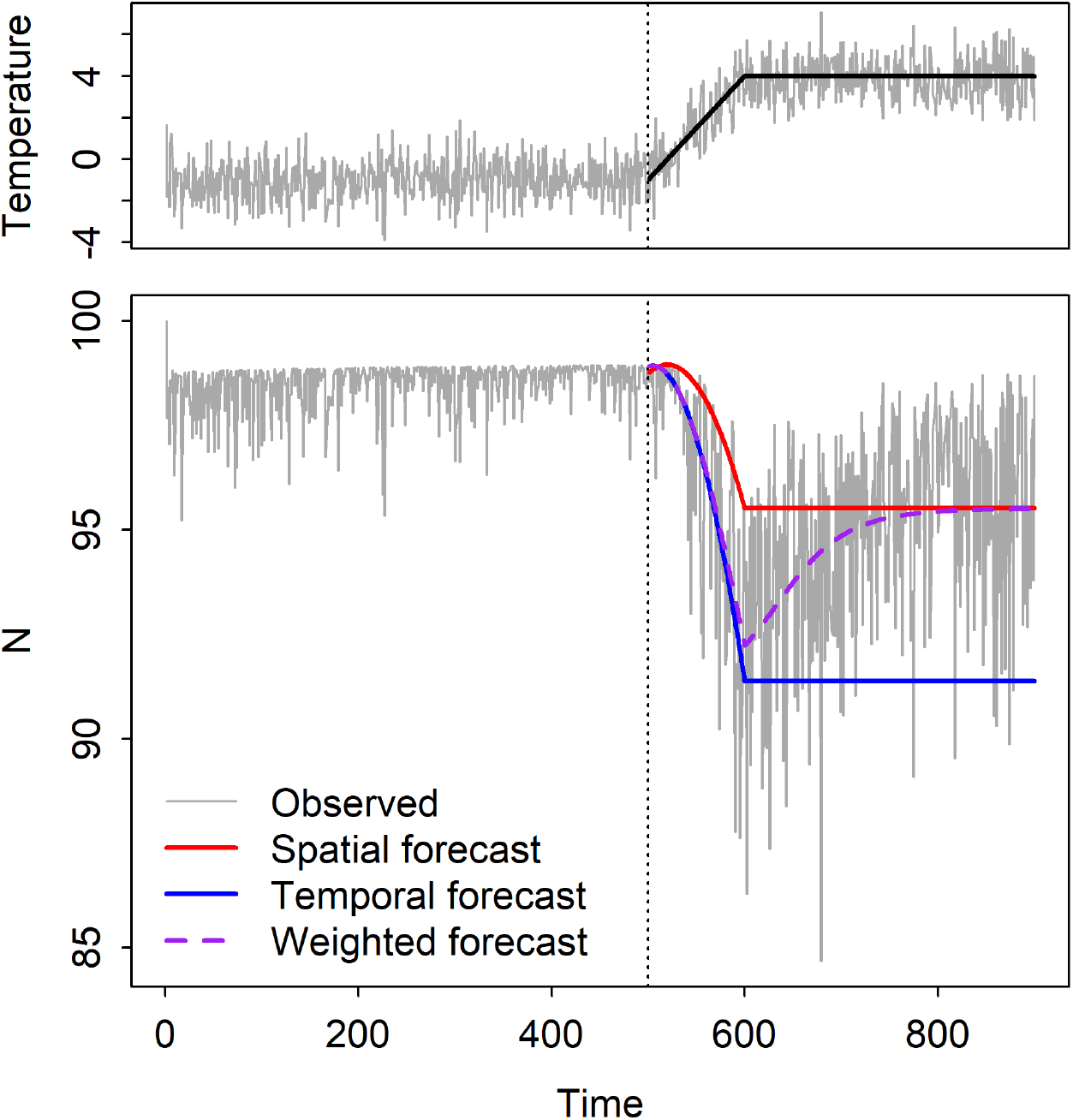
(Top) Simulated annual temperatures (grey) and expected temperature (black), which was used to make forecasts. (Bottom) Simulated population size and forecasts from the spatial, temporal and weighted statistical models.

With the onset of warming, the population crashed as the cold-adapted genotype decreased in abundance. Eventually, frequencies of the heterozygous genotype and the warm-adapted homozygous genotype began to increase and the population recovered (Fig. 5 bottom). The temporal statistical model (solid blue line in Fig. 5) accurately predicted the impact of the initial warming trend, but eventually became too pessimistic, while the spatial statistical model (solid red line in Fig. 5) did not handle the initial trend but accurately predicted the eventual, new steady state (Fig. 5 bottom).

As in the community turnover example, we also fit a weighted average of predictions from the spatial and temporal statistical models, with the weights changing over time. This weighted model initially reflected the temporal model (decrease from *t* = 500 to *t* = 600), but then rapidly transitioned to reflect the spatial model (*t* ≥ 700). The rapid transition in the weighting term, *ω*, occurred during the period of most rapid change in genotype frequencies (Fig. S-3). The weighted model’s predictions look impressively accurate, but, as in the community turnover example, that is because we used the full, simulated time series to fit the weighting term. A true forecast would require an independent method to predict how the model weights shift over time.

## Discussion

Ecological forecasts are typically made using either a space-for-time substitution approach based on models fit to spatial data or using dynamic models fit to time-series data. Our results demonstrate that these two approaches can make very different predictions about the future state of ecological systems. Which approach provides the most accurate forecasts depends on the forecast-horizon. In our simulations, time-series approaches performed best for short forecast horizons, whereas models based on spatial data made more accurate forecasts at long horizons. In addition, our simulations demonstrate extended transitional periods during which neither the time-series or the spatial approach was effective on its own. The challenge is determining what is “short-term,” what is “long-term,” and how to handle the many forecasts we need in ecology which fall in between. We have proposed that a weighted combination of the time-series and space-for-time approaches may produce better forecasts at these intermediate forecast horizons.

We designed our simulation studies to illustrate how the change in statistical model performance with increasing forecast horizon reflects differences in the types and scales of processes captured by spatial and temporal data sets. How could these hypotheses be tested with empirical data? The hypothesis that time-series models will be most effective for near-term forecasts already has empirical support, in the form of recent analyses of biodiversity forecasts at time ·, scales from one to ten years (Harris et al., 2018). The result should not be surprising, since local time-series data capture demographic processes, lagged effects, and responses of current assemblages to small changes in environmental conditions. In addition, the state of the system in the near future depends heavily on the current state. Since short-term forecasts do not typically require extrapolating into novel conditions, a model based on the historical range of variation which incorporates lags and accurate initial conditions is likely to be successful.

Space-for-time modeling approaches for predicting long-term, steady-state outcomes of ecological change have also been tested empirically, primarily via hind-casting. Overall, the results are mixed: some tests show reasonable prediction of changes in community composition (Blois et al., 2013; Illán et al., 2014) or species distributions (Norberg et al., 2019), supporting the hypothesis that datasets spanning spatial gradients capture the long-term outcome of interactions between fast processes and slower processes such as ecological and evolutionary selection, dispersal, and responses to large changes in the environment. Other attempts to validate predictions from space-for-time models have been discouraging (Worth et al., 2014; Illán et al., 2014; Davis et al., 2014; Brun et al., 2016; Veloz et al., 2012), indicating violations of model assumptions or effects of transient dynamics. However, predictions from the space-for-time approaches are rarely compared directly to predictions from time-series models (Harris et al. 2018 but see Renwick et al. 2018). We need more such comparisons to identify the appropriate modeling approach for different forecast horizons.

The greatest empirical challenge will be testing our hypothesis that a weighted average of spatial and temporal statistical models will make the best forecasts at intermediate time scales. There are two problems: finding appropriate data and determining the model weights *a priori*. Many data sets have both a longitudinal and spatial dimension, but we could not think of one which also featured a clear ecological response to significant environmental change. Surely such datasets exist, and we hope researchers who work with them will test our proposed weighted model. Determining model weights may be more difficult. In our simulations, we fit the weights directly to the simulated data, which is impossible to do for actual forecasting when the future is unknown. We need new theory or empirical case studies in order to assign these weights *a priori*.

Theory could explore the influence of different parameters on the rate at which slow processes begin to influence dynamics. The effects of some parameters are intuitive: in the community turnover example, increasing the fraction of dispersing individuals caused a more rapid shift in species composition and in model weights (Fig. 6A). Other parameters have less intuitive effects: we expected that increasing the temperature tolerance of genotypes in the evo-evolutionary example would accelerate the shift in model weights by maintaining higher genetic diversity. Our simulations showed the opposite effect, with wider tolerances slowing the shift in model weights (Fig. 6B), presumably by decreasing the strength of selection. Additional factors to consider include organism lifespans and the magnitude of directional environmental change relative to historical interannual variation.

**Figure 6:**
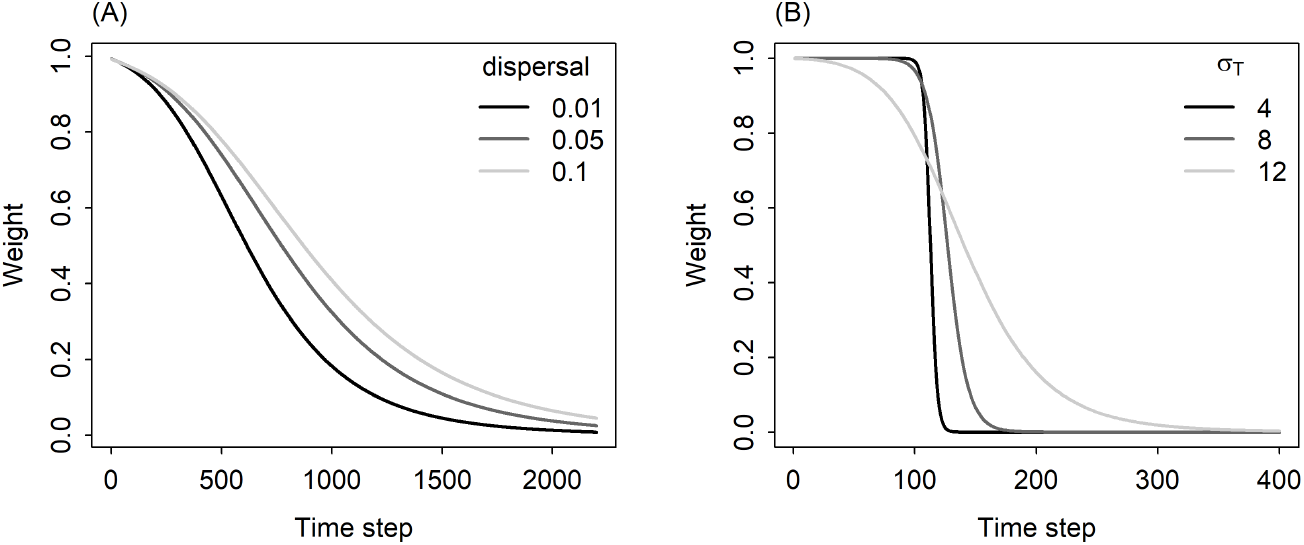
The rate of change in the weight of the temporal forecast (y-axis) depends on (A) the fraction of propagules dispersing in the community turnover example and (B) on the temperature tolerance of genotypes, given by *σ_T_* (larger values indicate wider thermal niches) in the eco-evolutionary example. Year 0 in these figures corresponds to the start of the temperature increase.

Empirical research could inform model weights by accumulating enough case studies to infer patterns in the weighting functions and guide applications in new systems. Developing rules of thumb would require testing many forecasts from both time-series and spatial models across a range of time-horizons. This effort may require a novel integration of typically disparate approaches, such as analyses of paleoecological data (e.g., Worth et al. 2014), long-term observational (e.g., Nice et al. 2019) or experimental data (e.g., Silvertown et al. 2006), and model systems with short-generation times (e.g., Good et al. 2017).

Given the challenges of determining model weights *a priori*, we should also pursue alternatives for intermediate forecast horizons. In the Introduction, we argued that fully process-based models are not feasible. However, a new class of statistical models offers a compromise between mechanistic detail and phenomenological feasibility. Spatiotemporal statistical modeling approaches are being developed to study patterns and processes of interest to ecological forecasters, such the spread of an invasive species or population status of a threatened species (Wikle, 2003; Williams et al., 2017; Schliep et al., 2018). Because these models include both fast processes, such as births and deaths, and slower processes, such as colonization and extinction dynamics, they have the potential to make better predictions at intermediate forecast horizons than purely spatial or temporal models. However, these spatiotemporal models have rarely been used in a forecasting context, due to a combination of data limitation and computational challenges. Many data sources contain either spatial or temporal variation, but not both, and when spatiotemporal datasets are available they often involve irregular sampling, creating challenges for modeling. Fitting and generating predictions from spatiotemporal models is also computationally intensive, especially with large datasets (McDermott and Wikle, 2017). Fortunately, thanks to large-scale monitoring efforts from remote sensing platforms, the National Ecological Observatory Network (https://www.neonscience.org/), and community science projects (e.g., eBird), large scale spatiotemporal data is increasingly available. In addition, new methods for spatiotemporal forecasting are being developed that address existing computational challenges (McDermott and Wikle, 2017), and access to high performance computing resources is increasingly common. Given these developments, future ecological forecasting efforts should explore spatiotemporal approaches and assess whether they improve predictions at intermediate time scales relative to traditional time-series or space-for-time approaches.

Our results have important implications for the emerging field of ecological forecasting. First, they suggest that evaluating model performance at both short and long forecsat horizons will be essential as research on forecasting methods accelerates. Second, while single approaches may perform reasonably well for either short or long horizons, skillful predictions at intermediate forecast horizons may require a combination of information from spatial and temporal statistical models. Intermediate time horizons pose challenges in other forecasting contexts as well. Weather forecasts based on regional-scale meteorological models are very effective for forecasting a week to ten days in advance, but then become largely uninformative. Forecasting these intermediate scales has been challenging in meteorology and will likely be challenging in ecology as well. While the recent emphasis on near-term iterative forecasting (Dietze et al., 2018) is the logical and tractable starting point, we also need to build understanding and capacity for forecasting ecological dynamics across all forecast horizons of interest.

### Data accessibility statement

The manuscript contains no original data. All computer code is available for reviewers in a zip archive that can be downloaded from Google Drive: https://drive.google.com/open?id=1aju04qtQvJmZG1mcpN-6hYBZILBhDfl2. Upon acceptance of the manuscript, code will be archived at Zenodo.

## Acknowledgements

We thank Heather Lynch and Juan Manuel Morales for comments that improved early drafts of the paper. PBA was supported by the National Science Foundation (DEB-1927282) and the Utah Agriculture Experiment Station (grant 1322). EPW was supported by the Gordon and Betty Moore Foundation’s Data-Driven Discovery Initiative (GBMF4563).

## Appendices

### A Spatial, temporal and spatial-temporal-weighted models

The two simulation models in the main text describe how population size, *N*(*x*, *t*), at location *x* changes over time (*t*). We assume that the temperature, *K*(*x*, *t*), at each location can vary in time and space. To forecast the dynamics generated by these simulations models, we fit a series of statistical models.

The spatial model, which we refer to as *S*, is a quadratic regression of the mean long-term population density at a location 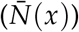 against the mean temperature at that location 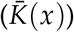. The quadratic term describes the unimodal relationship between 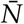 and 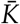. The spatial statistical model is

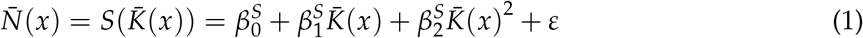

The temporal model, which we call *T*, starts with a time-series of “observed” population sizes, or total biomasses, at one location, *N*(*t*), for *t* = 1…*n* (the spatial index is suppressed because we only focus on one location at a time). In the community turnover example, we fit the following regression, which predicts biomass at time *t* + 1 as a function of biomass (*N*(*t*)) and annual temperature (*K*(*t*)) at time *t*,

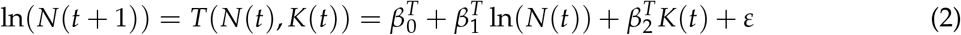

In the eco-evolutionary example, the response variable is the log of the population growth rate. The regression is

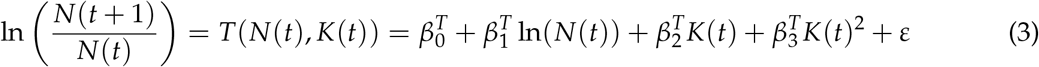

This version of the temporal model returns a per capita growth rate on the log scale. To predict population size at the next time step, we exponentiate the growth rate and multiply it by the current population size: exp(*T*(*N*(*t*), *K*(*t*)))*N*(*t*).

The weighted model is a weighted average of predictions from the spatial and temporal models, with the weights changing as a function of time, here expressed as the forecast horizon. The weights change as a function of the square root of the forecast horizon, to allow rapid shifts in the model weights.

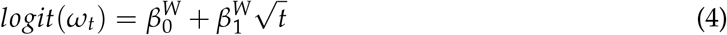

For the community turnover example, the predicted biomass from the weighted model is:

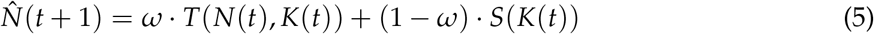

Again, we suppress the spatial subscript (*x*) here because we are focused on densities at just one location. For the eco-evolutionary example, the predicted population size from the weighted model is:

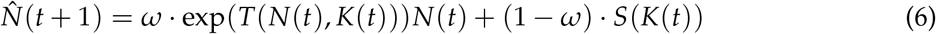

We used the `optim` function to estimate the *β^W^*s that minimize the sum of squared errors, 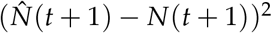.

In the main text, we show the point forecasts but not the uncertainty around the forecasts. After exploring that uncertainty, we decided that presenting it would be misleading. For the spatial and, especially, the temporal statistical models, the uncertainty is unrealistically low, because the models are estimated with very large samples sizes from the simulations. Furthermore, the simulations do not include noise; the only reason there is any uncertainty is because the statistical models are slightly mis-specified with respect to the process models. Showing uncertainty for the weighted model would be even less meaningful, because it is not a true, out-of-sample forecast (parameters are fit directly to the observations for which we make predictions). The R code to compute uncertainties for the spatial and temporal forecasts is available on our Github repository (https://github.com/pbadler/space-time-forecast), but is commented out.

### B Description of the meta-community model

Alexander et al. (2018) developed a meta-community model to represent dynamics of local communities arrayed along a one-dimensional elevation gradient, as influenced by three main processes: temperature-dependent growth, competition, and dispersal. Here we adapt their notation to be consistent our own.

The population size of species *i* in cell *x* at time *t* + 1, *N_i_* (*x*, *t* + 1), is computed in two steps. The first step accounts for changes in local population sizes due to dispersal. In each local community, all species export a fraction (*d*) of their local population to the two adjacent communities in the 1-dimensional landscape:

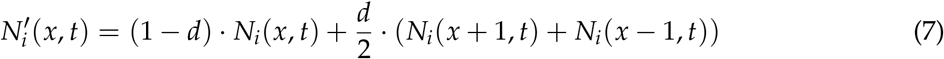

Here *N*′ distinguishes the post-dispersal population size from the pre-dispersal population size. The second step computes population growth, taking into account competition:

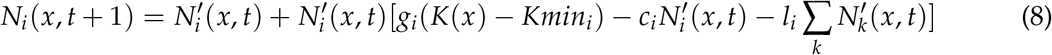

In the absence of competition, the growth rate (*g_i_*) is determined by the difference between the temperature at site *x* (*K*(*x*)) and the focal species’ minimum temperature tolerance, *Kmin_i_*, the lowest temperature at which a species can maintain a positive growth rate. Growth is further reduced by intraspecific and interspecific competition, parameterized by *c_i_* and *l_i_*. All species are assigned the same value of *c_i_*, which represents an additional effect of intraspecific competition on top of interspecific competition. This stabilizes coexistence, since every species will exert stronger intra- than interspecific competition. However, values of *l* vary among species to create a trade-off between growth rates and competitive ability versus low temperature tolerance: fast-growing species (high *g_i_*) are more tolerant of interspecific competition (low *l_i_*) but are more limited by temperature (high *Kmin_i_*).

### C Description of the eco-evolutionary annual plant model

#### Haploid Model

Begin with a haploid model that describes the number of seeds present in a seed bank. *N_i,t_* is the number of seeds of species *i* at time *t*. The model is

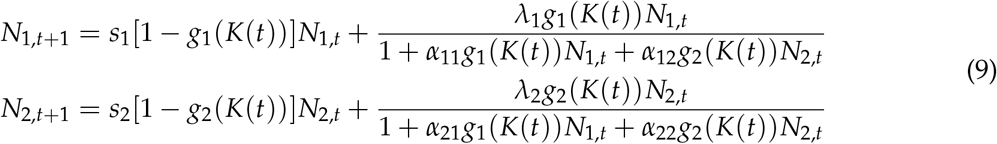

where *g_i_*(*K*(*t*)) is the probability of germination, *K*(*t*) is the temperature at time *t*, *s_i_* is the seed survival probability for species *i*, and *λ_i_* is the seed production rate per plant. Below we refer to the *α_ij_* as intra- and inter-genotype competition coefficients.

#### Diploid Model

Consider a one-species diploid model. The genotypes are denoted by *AA*, *Aa*, and *aa*. The number of each genotypes at time *t* is *N_AA_*(*t*), *N_Aa_*(*t*), and *N_aa_*(*t*). The germination rates for each genotype are *g_AA_*(*K*(*t*)), *g_Aa_*(*K*(*t*)), and *g_aa_*(*K*(*t*)). The seed survival probability and seed production rate for genotype *AA* are *s_AA_* and *λ_AA_*, respectively. The analogous parameters for the other genotypes are similarly denoted. The competition coefficients are denoted by *α_i,j_*, e.g., *α_AA,AA_* or *α_AA,Aa_*. Throughout we assume that gametes mix randomly in the population.

First consider the case where the competition coefficients are zero (*α_i,j_* = 0). Let *T* denote the total number of gamete-pairs produced in a given year,

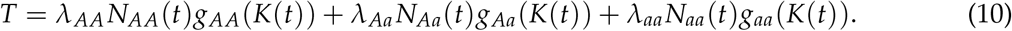

The first term is the number of gamete-pairs produced by *AA* individuals. The second and third terms are the numbers of gamete-pairs produced by *Aa* and *aa* individuals, respectively. The proportion of *A* gametes (*ϕ_A_*) and the proportion of *a* gametes (*ϕ_a_*) are given by

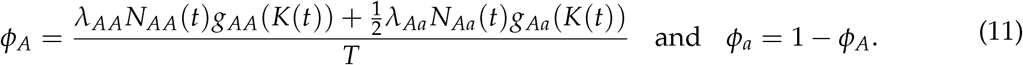

Note that the *T* in the denominator of *ϕ_A_* shows up because we are computing proportions. Combining all of these we get the dynamics for each genotype,

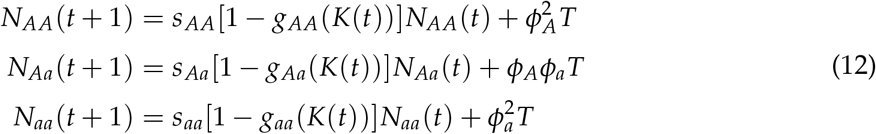

Now consider the case where the competition coefficients are non-zero (*α_i,j_* ≠ 0). Including competition changes the way in which we compute *T*, *ϕ_A_*, and *ϕ_a_*. Specifically, because the total number of seeds produced per year by each genotypes is reduced based on intra- and intergenotype competition, the total number of gamete-pairs becomes

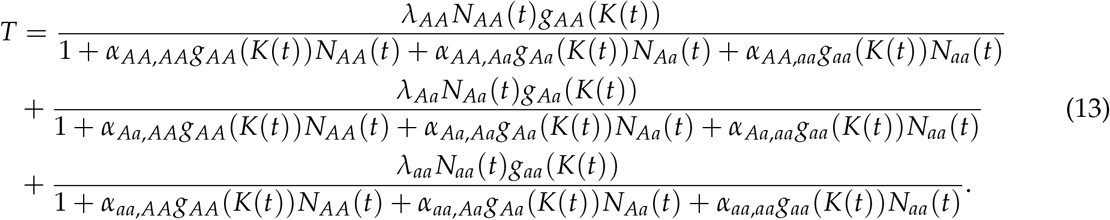

The first line is the number of gamete-pairs produced by *AA* individuals after accounting for the effects of competition. The second and third lines are the numbers of gamete-pairs produced by *Aa* and *aa* individuals, respectively. The proportions of *A* gametes and *a* gametes are

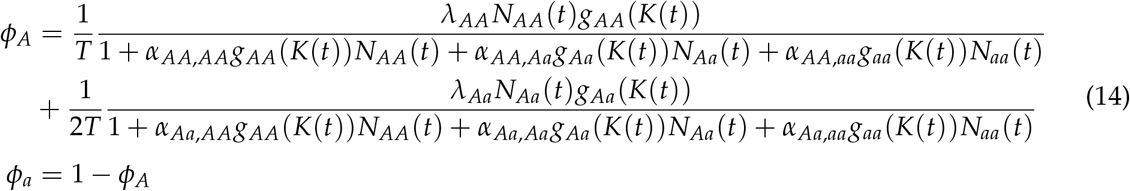

Combining all of this results in the same model as above,

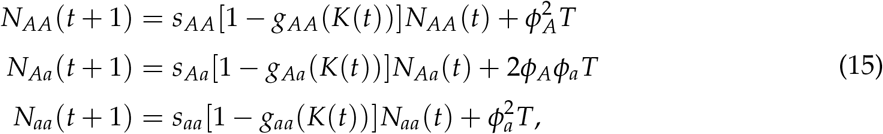

but the definitions of *T*, *ϕ_A_*, and *ϕ_a_* are given by equations (13) and (14).

### D Supplementary Figures

**Figure S-1:**
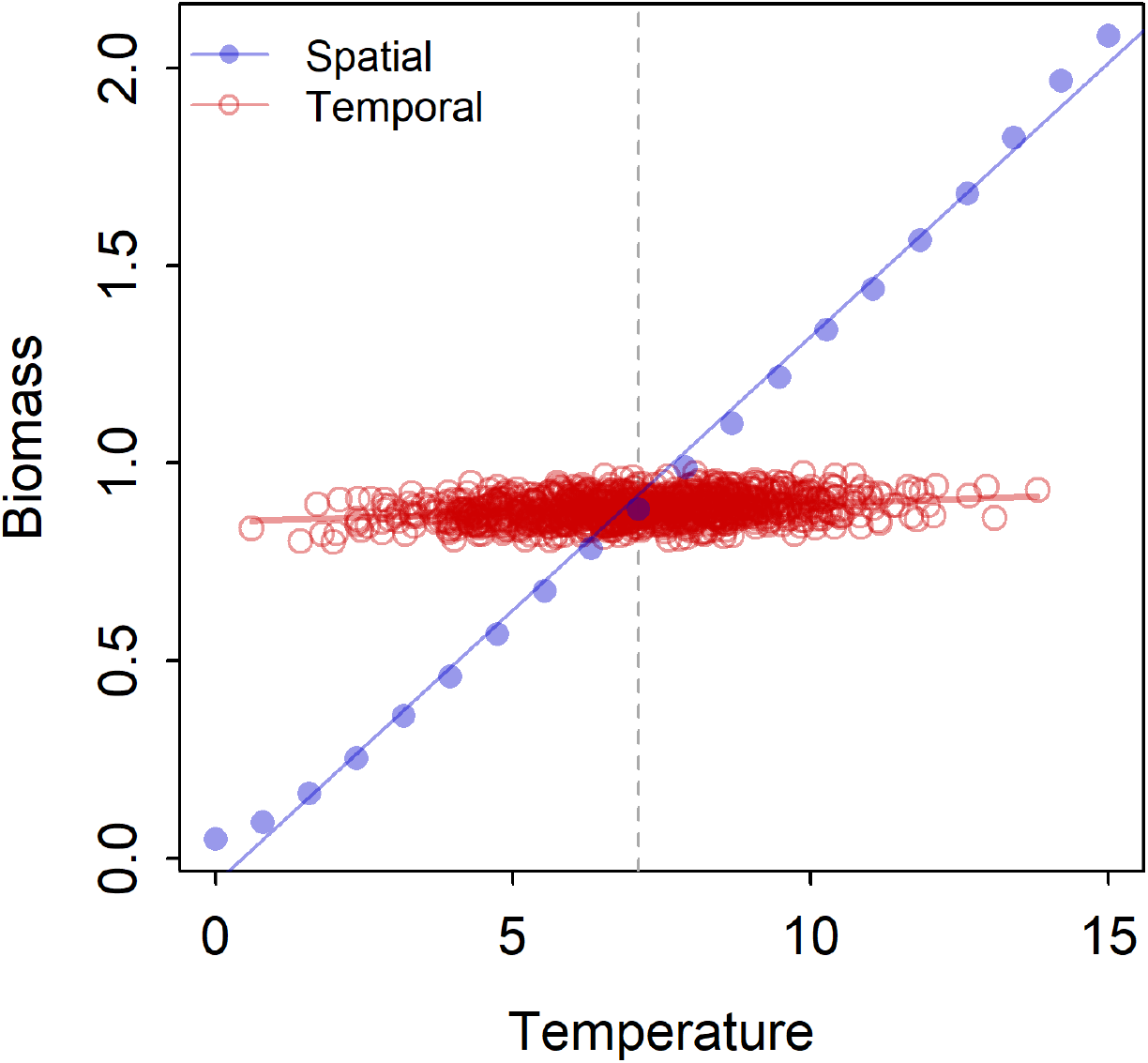
(Results for total biomass from the community turnover model. Blue points show mean total biomass during the baseline period at locations across the temperature gradient, and the blue line shows predictions from the spatial model. Red points show annual total biomass during the baseline period as a function of annual temperature at the central site on the gradient. The red line shows predictions from the temporal model.

**Figure S-2:**
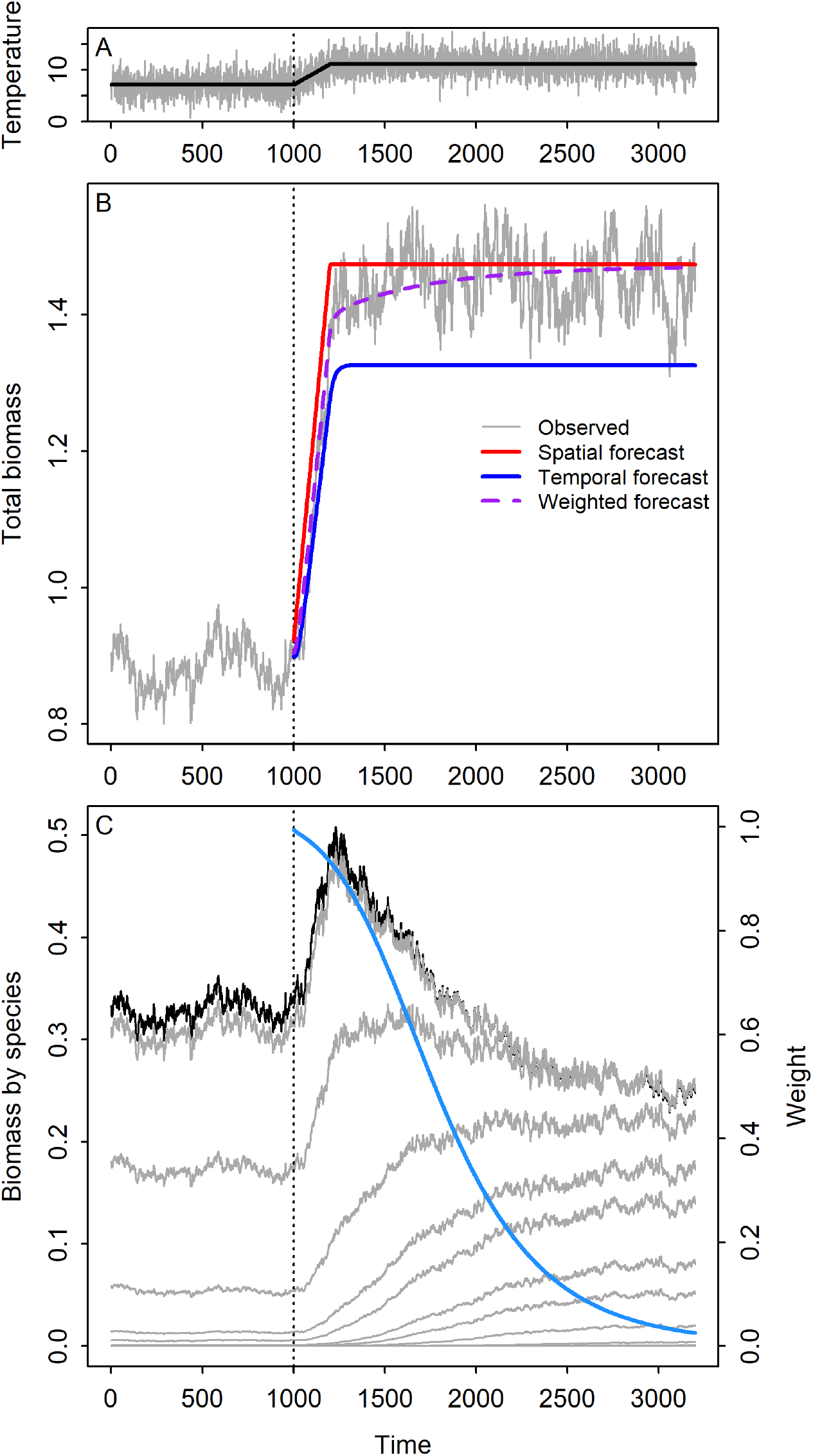
Results for total biomass from the community turnover model. (A) Simulated annual temperatures (grey) and expected temperature (black), which was used to make forecasts, at the focal site. (B) Simulated total biomass and forecasts from the spatial, temporal and weighted models. (C) Simulated changes in biomass of all species (grey) at the focal site in the metacommunity model, and the weight given to the temporal model for total biomass (blue). Year 1000 in this figure corresponds to the start of the temperature increase.

**Figure S-3:**
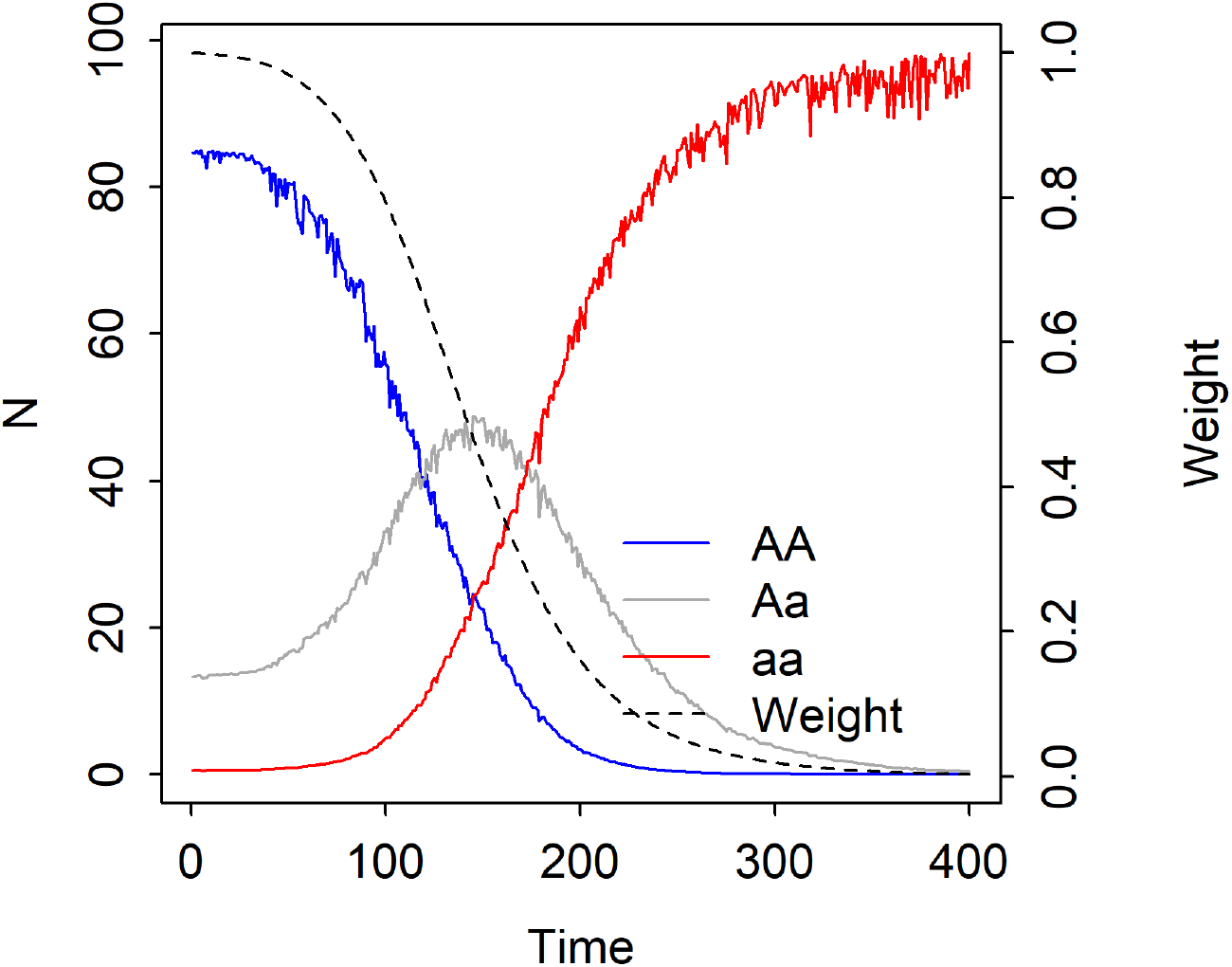
Simulated shifts in genotype abundances, and the model weighting term, *ω*, during the warming phase and the following stationary temperature phase. Year 0 in this figure corresponds to the start of the temperature increase.

## Literature cited

Alexander, J. M. et al. 2018. Lags in the response of mountain plant communities to climate change. – Global Change Biology 24: 563–579.

Blois, J. L. et al. 2013. Space can substitute for time in predicting climate-change effects on biodiversity. – Proceedings of the National Academy of Sciences 110: 9374–9379.

Brun, P. et al. 2016. The predictive skill of species distribution models for plankton in a changing climate. – Global Change Biology 22: 3170–3181.

Chevalier, M. and Knape, J. 2020. The cost of complexity in forecasts of population abundances is reduced but not eliminated by borrowing information across space using a hierarchical 9 approach. – Oikos in press.

Clark, J. et al. 2001. Ecological forecasts: An emerging imperative. – Science 293: 657–660.

Dalgleish, H. J. et al. 2011. Climate influences the demography of three dominant sagebrush steppe plants. – Ecology 92: 75–85.

Davis, E. B. et al. 2014. Ecological niche models of mammalian glacial refugia show consistent bias. – Ecography 37: 1133–1138.

Dietze, M. C. 2017. Prediction in ecology: a first-principles framework. – Ecological Applications 27: 2048–2060.

Dietze, M. C. et al. 2018. Iterative near-term ecological forecasting: Needs, opportunities, and challenges. – Proceedings of the National Academy of Sciences 115: 1424–1432.

Elith, J. and Leathwick, J. R. 2009. Species distribution models: ecological explanation and prediction across space and time. – Annual Review of Ecology, Evolution, and Systematics 40: 677–697.

Good, B. H. et al. 2017. The dynamics of molecular evolution over 60,000 generations. – Nature 551: 45.

Harris, D. J. et al. 2018. Forecasting biodiversity in breeding birds using best practices. – PeerJ 6: e4278.

Hefley, T. J. et al. 2017. When mechanism matters: Bayesian forecasting using models of ecological diffusion. – Ecology Letters 20: 640–650.

Hoffmann, A. A. and Sgro, C. M. 2011. Climate change and evolutionary adaptation. – Nature 470: 479–485.

Houlahan, J. E. et al. 2017. The priority of prediction in ecological understanding. – Oikos 126: 1–7.

Hyndman, R. and Athanasopoulos, G. 2018. Forecasting: Principles and Practice. – OTexts.

Illán, J. G. et al. 2014. Precipitation and winter temperature predict long-term range-scale abundance changes in Western North American birds. – Global Change Biology 20: 3351–3364.

Kleinhesselink, A. R. and Adler, P. B. 2018. The response of big sagebrush (Artemisia tridentata) to interannual climate variation changes across its range. – Ecology 99: 1139–1149.

Lauenroth, W. K. and Sala, O. E. 1992. Long-term forage production of North American short-grass steppe. – Ecological Applications 2: 397–403.

Levin, S. A. 1992. The problem of pattern and scale in ecology: the robert h. macarthur award lecture. – Ecology 73: 1943–1967.

McDermott, P. L. and Wikle, C. K. 2017. An ensemble quadratic echo state network for non-linear spatio-temporal forecasting. – Stat 6: 315–330.

Mouquet, N. et al. 2015. REVIEW: Predictive ecology in a changing world. – Journal of Applied Ecology 52: 1293–1310.

Nice, C. C. et al. 2019. Extreme heterogeneity of population response to climatic variation and the limits of prediction. – Global Change Biology 25: 2127–2136.

Norberg, A. et al. 2019. A comprehensive evaluation of predictive performance of 33 species distribution models at species and community levels. – Ecological Monographs 89: e01370.

Renwick, K. M. et al. 2018. Multi-model comparison highlights consistency in predicted effect of warming on a semi-arid shrub. – Global Change Biology 24: 424–438.

Rosenzweig, M. L. et al. 1995. Species diversity in space and time. – Cambridge University Press.

Sala, O. E. et al. 1988. Primary production of the central grassland region of the United States. – Ecology 69: 40–45.

Schliep, E. M. et al. 2018. Joint species distribution modelling for spatio-temporal occurrence and ordinal abundance data. – Global Ecology and Biogeography 27: 142–155.

Silvertown, J. et al. 2006. The Park Grass Experiment 1856-2006: Its contribution to ecology. – Journal of Ecology 94: 814.

Urban, M. C. et al. 2012. On a collision course: competition and dispersal differences create no-analogue communities and cause extinctions during climate change. – Proceedings of the Royal Society of London B: Biological Sciences 279: 2072–2080.

Veloz, S. D. et al. 2012. No-analog climates and shifting realized niches during the late quaternary: implications for 21st-century predictions by species distribution models. – Global Change Biology 18: 1698–1713.

Ward, E. J. et al. 2014. Complexity is costly: a meta-analysis of parametric and non-parametric methods for short-term population forecasting. – Oikos 123: 652–661.

Wikle, C. K. 2003. Hierarchical Bayesian Models for Predicting the Spread of Ecological Processes. – Ecology 84: 1382–1394.

Williams, J. W. and Jackson, S. T. 2007. Novel climates, no-analog communities, and ecological surprises. – Frontiers in Ecology and the Environment 5: 475–482.

Williams, P. J. et al. 2017. An integrated data model to estimate spatiotemporal occupancy, abundance, and colonization dynamics. – Ecology 98: 328–336.

Worth, J. R. P. et al. 2014. Environmental niche modelling fails to predict Last Glacial Maximum refugia: niche shifts, microrefugia or incorrect palaeoclimate estimates? – Global Ecology and Biogeography 23: 1186–1197.

